# Systemic analysis of tissue cells potentially vulnerable to SARS-CoV-2 infection by the protein-proofed single-cell RNA profiling of ACE2, TMPRSS2 and Furin proteases

**DOI:** 10.1101/2020.04.06.028522

**Authors:** Lulin Zhou, Zubiao Niu, Xiaoyi Jiang, Zhengrong Zhang, You Zheng, Zhongyi Wang, Yichao Zhu, Lihua Gao, Hongyan Huang, Xiaoning Wang, Qiang Sun

**Affiliations:** Institute of Biotechnology, 20 Dongda Street, Beijing 100071, P.R. China; School of Medicine, Nankai University, 94 Weijin Road, Tianjin, 300071, P. R. China; Department of Oncology, Beijing Shijitan Hospital of Capital Medical University, 10 TIEYI Road, Beijing 100038, P. R. China; School of Laboratory Medicine and Biotechnology, Southern Medical University, Guangzhou 510515, P. R. China; National Clinic Center of Geriatric, the Chinese PLA General Hospital, Beijing 100853, P. R. China

**Keywords:** COVID-19, SARS-CoV-2, ACE2, TMPRSS2, Furin, single cell RNA profiling, pscRNA profiling

## Abstract

Single-cell RNA profiling of ACE2, the SARS-CoV-2 receptor, had proposed multiple tissue cells as the potential targets of SARS-CoV-2, the novel coronavirus causing the COVID-19 pandemic. However, most were not echoed by the patients’ clinical manifestations, largely due to the lack of protein expression information of ACE2 and co-factors. Here, we incorporated the protein information to analyse the expression of ACE2, together with TMPRSS2 and Furin, two proteases assisting SARS-CoV-2 infection, at single cell level *in situ*, which we called protein-proofed single-cell RNA (pscRNA) profiling. Systemic analysis across 36 tissues revealed a rank list of candidate cells potentially vulnerable to SARS-CoV-2. The top targets are lung AT2 cells and macrophages, then cardiomyocytes and adrenal gland stromal cells, followed by stromal cells in testis, ovary and thyroid. Whereas, the polarized kidney proximal tubule cells, liver cholangiocytes and intestinal enterocytes are less likely to be the primary SARS-CoV-2 targets as ACE2 localizes at the apical region of cells, where the viruses may not readily reach. Actually, the stomach may constitute a physical barrier against SARS-CoV-2 as the acidic environment in normal stomach (pH < 2.0) could completely inactivate SARS-CoV-2 pseudo-viruses. These findings are in concert with the clinical characteristics of prominent lung symptoms, frequent heart injury, and uncommon intestinal symptoms and acute kidney injury. Together, we provide a comprehensive view on the potential SARS-CoV-2 targets by pscRNA profiling, and propose that, in addition to acute respiratory distress syndrome, attentions should also be paid to the potential injuries in cardiovascular, endocrine and reproductive systems during the treatment of COVID-19 patients.

## INTRODUCTION

In January, 2020, a novel coronavirus of unknown origin was identified to cause severe pneumonia in about 15-20% infected patients (Guan et al., 2020; Novel Coronavirus Pneumonia Emergency Response Epidemiology, 2020), the disease is currently called COVID-19 abbreviated from Coronavirus Disease 2019 (Zhu et al., 2020a). The virus is phylogenetically similar (~76% amino acid identity) to the severe acute respiratory syndrome coronavirus (SARS-CoV) (Tang et al., 2020; Wu et al., 2020b; Xu et al., 2020; Zhou et al., 2020b; Zhu et al., 2020b), and was subsequently named as SARS-CoV-2 after the initial name of 2019-nCoV. The SARS-CoV-2 virus is much more contagious than SARS-CoV, and had infected more than 12 million of individuals from 215 countries and territories as of July 12, 2020, leading to more than 56 thousands of deaths with an average motility rate of about 4.5% (WHO, 2020). The pandemic of COVID-19 is posing a global health emergency.

The coronavirus is a large group of enveloped, single-strand positive-sense RNA viruses, with SARS-CoV and Middle East respiratory syndrome coronavirus (MERS-CoV) the two known deadly viruses for human (Li, 2016). The spike (S) envelope glycoproteins on coronavirus are the major determinants of host cell entry. Proteolytic cleavage of S protein produces S1, the N-terminal region of S protein that is responsible for receptor binding, and S2, the trans-membrane C-terminal region of S protein that promotes membrane fusion. The cleavage step is often permissive for the fusion function of S protein as it helps to release the fusion peptide to insert into the target cellular membrane (Li, 2016; Millet and Whittaker, 2015). Therefore, the host range and cell/tissue tropism of coronaviruses were believed to be controlled by the S protein engagement of host cell receptor, and by the proteolytic cleavage of the S protein as well (Millet and Whittaker, 2015). Recently, works from several groups demonstrated, either bioinformatically or experimentally, that angiotensin-converting enzyme 2 (ACE2), the receptor for SARS-CoV virus (Li et al., 2003), is also a functional cellular receptor for SARS-CoV-2 virus (Hoffmann et al., 2020; Walls et al., 2020; Wrapp et al., 2020; Xu et al., 2020; Zhou et al., 2020b), and transmembrane protease serine 2 (TMPRSS2) and Furin are two proteases that process SARS-CoV-2 S protein to establish efficient infection (Hoffmann et al., 2020; Li et al., 2020; Meng et al., 2020; Walls et al., 2020).

Single-cell RNA (scRNA) profiling is a state-of-the-art tool to dissect gene expression at single cell level, therefore was employed to explore the target cells of the SARS-CoV-2. Based on the profiling of ACE2 mRNA expression in different tissues/organs, multiple types of cells were proposed to be potentially targeted by SARS-CoV-2 virus, including the lung alveolar type 2 (AT2) cells (Zhao et al., 2020a), nasal epithelial cells (WU et al., 2020a), oesophageal epithelial cells and intestinal enterocytes (Zhang et al., 2020), liver cholangiocytes (Chai et al., 2020), cardiomyocytes (Xin Zou), kidney proximal tubule cells (Lin et al., 2020), spermatogonia and Leydig/sertoli cells in the testis (Zhengpin and Xiaojiang, 2020). However, unparallel to the many potential targets cells and organs proposed, the COVID-19 patients primarily displayed typical symptoms of inflammation in the lung, where only a very small portion of cells (~0.64% of total cells, and ~1.4% of AT2 cells) expressed ACE2 mRNA (Zhao et al., 2020a); meanwhile, the injuries in other organs/tissues, such as the kidney and the intestinal track, where ACE2 gene was expressed at high levels, seemed to be uncommon (Guan et al., 2020; Wang et al., 2020a). This obvious discrepancy suggests that mechanisms other than ACE2 mRNA levels are also involved in the regulation of SARS-CoV-2 infection of their target cells.

Considering that mRNA level does not always dictate comparable protein expression and subcellular localizations, which are missing information from mRNA profiling, are critical for protein functions, we set out to explore ACE2 expression at both mRNA and protein levels by taking advantages of the curated public database. We called this method as protein-proofed scRNA (pscRNA) profiling. Moreover, we also analysed the co-expression of ACE2 with its two processing proteases, TMPRSS2 and Furin, at single cell resolution *in situ* by pscRNA profiling. Systemic analysis of 36 human tissues/organs revealed that 1) a rank list of potential SARS-CoV-2 targets with lung AT2 cell and macrophages as the top targets, then cardiomyocytes, and stromal cells in testis, ovary, adrenal and thyroid glands. Among them, the lung macrophages, stromal cells in ovary and adrenal gland, were identified for the first time, which may account for severe clinical symptoms and rapid disease progression; 2) the mRNA levels may differ dramatically from the protein levels for ACE2, TMPRSS2 and Furin in different tissue cells; and protein subcellular localization is another factor potentially affecting virus host-entry; 3) the co-expression of ACE2 with TMPRSS2 and Furin proteases may contribute to establish efficient infection of SARS-CoV-2 virus.

## MATERIAL AND METHODS

### Analysis Framework

To achieve a comprehensive analysis of tissue cells potentially vulnerable to SARS-CoV-2 virus, we employed a step-in strategy, ie, from tissue to cell, from multiple cells to single cell, from protein to mRNA, from single gene expression to co-expression. During analysis, we primarily focused on the expression of ACE2 while taking into account its co-expression with TMPRSS2 and Furin, two proteases that were believed to facilitate SARS-CoV-2 infection. To evaluate the cell vulnerability, not only the mRNA levels but also the protein levels were considered, and the protein levels actually take more weights as protein is the main function executor; also, not only protein levels but also their subcellular localizations in a specified type of cell were considered, as the subcellular localization determines the routes whereby viruses might access the protein receptor, for instance, apical localized surface protein would primarily be accessed by viruses from the luminal side, but not viruses from blood stream that are more likely infected unpolarized stromal cells. By following above principles, we firstly examined tissue distribution of ACE2, TMPRSS2 and Furin in both RNA and protein levels, then analyzed their expressions in situ by immunohistochemistry, which could provide information on both protein levels and subcellular localization. Subsequently, single cell RNA profiling was performed to determine and confirm cell type and co-expression pattern. Finally, a rank list was proposed by integrating information from RNA and protein levels, protein subcellular localizations, cell types and co-expression pattern, and the available experimental evidences and clinical manifestations.

### Data Acquisition

For scRNA profiling, the raw gene expression matrices for single cells were downloaded from Gene Expression Omnibus (GEO) database(https://www.ncbi.nlm.nih.gov/geo/) and Human Cell Atlas Data Portal (https://data.humancellatlas.org/). In total, we acquired the single cell gene expression datasets of various normal tissues and organs, including the lung, heart, liver, kidney, intestine, stomach, testis, ovary, adrenal gland and thyroid. Apart from the heart dataset, which was obtained by the smart-seq2 method, a majority of the single cell data were generated through the 10x Chromium platform. The GEO accession number of these datasets is GSE122960 (lung) (Reyfman et al., 2019), GSE109816 (heart) (Wang et al., 2020b), GSE131685 (kidney) (Liao et al., 2020), GSE125970 (intestine) (Wang et al., 2020c), GSE134520 (stomach) (Zhang et al., 2019a), GSE109037 (testis) (Hermann et al., 2018), GSE134355 (thyroid and adrenal gland)(Han et al., 2020), and GSE118127 (ovary) (Fan et al., 2019). The liver dataset was acquired from the Human Cell Atlas Data Portal (MacParland et al., 2018). The summary of sample information for single-cell data was shown in the supplementary table 1.

For tissue distribution of mRNA and protein expression profiles, data were obtained for target genes from the “TISSUE” categories of “THE HUMAN PROTEIN ATLAS” (http://www.proteinatlas.org/) (Uhlen et al., 2015). The mRNA expression from the Genotype-Tissue Expression (GTEx) database was chosen for demonstration with reference to the normalized consensus dataset (Carithers and Moore, 2015). The protein expression scores were extracted directly from the “PROTEIN EXPRESSION SUMMARY” subcategory from the Human Protein Atlas (HPA) database. Immunohistochemistry (IHC) staining was performed on normal human tissue samples (Thul et al., 2017). ACE2 expression was primarily detected with the rabbit antibody (1:250, HPA000288, Sigma-Aldrich) and confirmed with the mouse antibody (1:5000, CAB026174, R&D Systems); TMPRSS2 expression was detected with the rabbit antibody (1:300, HPA035787, Sigma-Aldrich); Furin expression was detected with the rabbit antibody (1:125, sc-20801, Santa Cruz Biotechnology). The information on expression intensity and cell types expressing respective genes was extracted from the staining reports for each staining in the HPA database.

### Quality Control and Data Normalization

The raw count matrices of single-cell transcriptome were imported into R (version 3.6.1, https://www.r-project.org/) and processed by the Seurat R package(version3.1.4) (Stuart et al., 2019). The filter criteria for low-quality cells were determined on the basis of the number of genes and the percentage of mitochondrial genes in the distinct tissue samples. Generally, cells with less than 200 detected genes, higher than 2500 detected genes and higher than 25% mitochondrial genome transcript ratio were removed. In specific, cells with higher than 72% mitochondrial genome transcript ratio in the heart dataset and cells with higher than 50% mitochondrial genome transcript ratio in the liver dataset were removed as mentioned by the corresponding authors (MacParland et al., 2018; Wang et al., 2020b). Then, the gene expression matrices were normalized and scaled. Briefly, for each cell, the expression counts for each gene were divided by the sum of counts for all genes of that cell, multiplied by a scaling factor (10,000) and log transformed using the “NormalizeData” function in the R package Seurat. Furthermore, 2000 highly variable genes were selected based on a variance stabilizing transformation method for downstream analysis.

### Data Integration and Dimension Reduction

To avoid batch effects among samples and experiments in the heart and intestine datasets, we adopted the anchoring procedure in the Seurat R package to integrate the datasets as described previously (Stuart et al., 2019). In brief, the “anchor” correspondences between datasets were identified using the “FindIntegrationAnchors” function with default parameters in Seurat. Then, we used the “IntegrateData” function in Seurat to obtain the batch-corrected expression matrices. We also applied another integration approach called “Harmony” (Korsunsky et al., 2019) to the kidney and testis datasets. The Harmony R package (version1.0, https://github.com/immunogenomics/Harmony) focuses on scalable integration of the scRNA-seq data for batch correction.

In order to reduce the number of dimensions representing each cell, the “RunPCA” function in Seurat was performed to calculate principal components (PCs). Then, the number of PCs used in downstream analysis and visualization was determined based on the “JackStraw” procedure and the elbow of a scree plot. Nonlinear dimensionality reduction algorithms, including uniform manifold approximation and projection (UMAP) and t-distributed stochastic neighbour embedding (t-SNE), were also used to conduct unsupervised clustering of single cells. Specifically, we made use of the UMAP and t-SNE to place cells with similar local neighbourhoods based on the statistically significant PCs to visualize the datasets.

### Cell Clustering, Annotation and Visualization

We applied a graph-based clustering approach to assemble cells. Briefly, we constructed a K-nearest neighbour graph based on the Euclidean distance in principal component analysis(PCA) space and refined the edge weights between any two cells based on the shared overlap in their local neighbourhoods through the “FindNeighbors” function in Seurat. Subsequently, we applied the Louvain algorithm using the “FindCluster” function in Seurat to group cells.

We applied two strategies to determine cell types. First, canonical marker genes were applied to annotate each cluster into known biological cell types. Expression dot plots or heatmaps of the marker genes for each cluster are shown in Figure 3. Second, we used the “FindAllMarkers” function from R package Seurat to find differentially expressed genes by comparing each cluster of cells with all other cells. Differentially expressed genes in each cluster were enriched to the cell markers gene sets(Zhang et al., 2019b) using the “clusterProfiler” R package (version 3.14.3) (Yu et al., 2012) to predict the probable cell types.

Umap plots, heatmaps, violin plots, and dot plots were generated using the R package Seurat. In order to visualize co-expression of two genes simultaneously, we adopted the “FeaturePlot” function and “FeatureScatter” function in Seurat.

### Cell culture, constructs and pseudovirus production

The HEK293T, 293T-ACE2 and Hela-ACE2 cells were maintained in DMEM (MACGENE Tech Ltd., Beijing, China) supplemented with 10% fetal bovine serum (Kang Yuan Biol, Tianjin, China) and 1% Penicillin-Streptomycin (MACGENE Tech Ltd., Beijing, China). All cells were incubated with 5% CO2 at 37°C. The codon-optimized SARS-CoV-2 S cDNA was synthesized at Genscript Biotech Corporation (Nanjing, China). The wild type S genes of SARS-CoV-2 were cloned into pSecTag2-Hygro-A through seamless homologous recombination.

For the production of MSV-based SARS-CoV-2 pseudotypes, HEK293T cells were co-transfected with an S encoding-plasmid, a Gag-Pol packaging construct (Addgene, 8449, USA) and the pQCXIP retroviral vector (Clontech, USA) expressing a luciferase reporter by using Lipofectamine LTX and Plus Reagent (Invitrogen, 1784283, USA) according to the manufacturer’s instructions. Cells were incubated for 6 hours at 37 °C in transfection medium, then DMEM containing 10% FBS and cultured for additional 48 hours. The supernatants were harvested and filtered through 0.45 μm membranes and stored at −80 °C.

### Pseudovirus assay

Before infection, a defined amount of supernatant containing pseudo-viruses were mixed with certain volumes of hydrochloric acid stock buffer (0.5 M) to produce virus fluids of different pH (1, 2, 4 or 7, respectively). The mixture experiments were performed in triplicates, which gave consistent final pH values with a variation range within 0.1 pH units. Virus mixtures were incubated at room temperature for 15 minutes, after which they were neutralized by 1 M NaOH solution and laid at room temperature for additional 15 minutes. All pH values were determined by a PH meter (FiveEasy™ plus, Mettler Toledo, Switzerland). Please find detail recipe for pH calibration in Supplementary Table 3.

293T-ACE2 cells and Hela-ACE2 cells were cultured in DMEM medium supplemented with 10% FBS and 1% PenStrep. The viral fluids were mixed with normal culture media containing 0.5 × 10^4^ cells and plated into a 96-well plate with a final volume of 100 ul per well. At the time points of 24 hours, or 48 hours post infection, 100 μL One-Glo-EX (Promega, E6120) was added into the cells for incubation in the dark for 10 min prior to reading on an Enspire 2300 multilable reader (Perkin Elmer, USA). Measurements were done at least in triplicates and relative luciferase units (RLU) were plotted.

## RESULTS

### Tissue distribution of ACE2, TMPRSS2 and Furin proteases

Based on the expression analysis across 36 human tissues, ACE2 displayed a tissue-specific expression at both mRNA and protein levels. A total of 10 tissues expressed relatively higher level of ACE2 mRNA, including the small intestine, colon, oesophagus, thyroid gland, kidney, testis, ovary, breast, heart muscle and adipose tissue. Meanwhile, the majority of the other tissues, such as the lung, liver, pancreas, spleen, prostate, urinary bladder, skin and neuronal tissues, had the marginal expression of ACE2 mRNA (Figure 1A-B, S1A). The protein expression score, though also displaying a tissue-specific pattern, only indicated that 6 tissues expressed ACE2 protein, with 5 of them matched the mRNA expression, including the small intestine, colon, thyroid gland, kidney and testis. Interestingly, while the adrenal gland expressed little mRNA, it had a median level of ACE2 protein expression. This inconsistency was true for ACE2 to other tissues such as the breast, heart muscle and adipose tissue, which expressed high levels of ACE2 mRNA but had undetectable levels of ACE2 protein.

**Figure 1.**
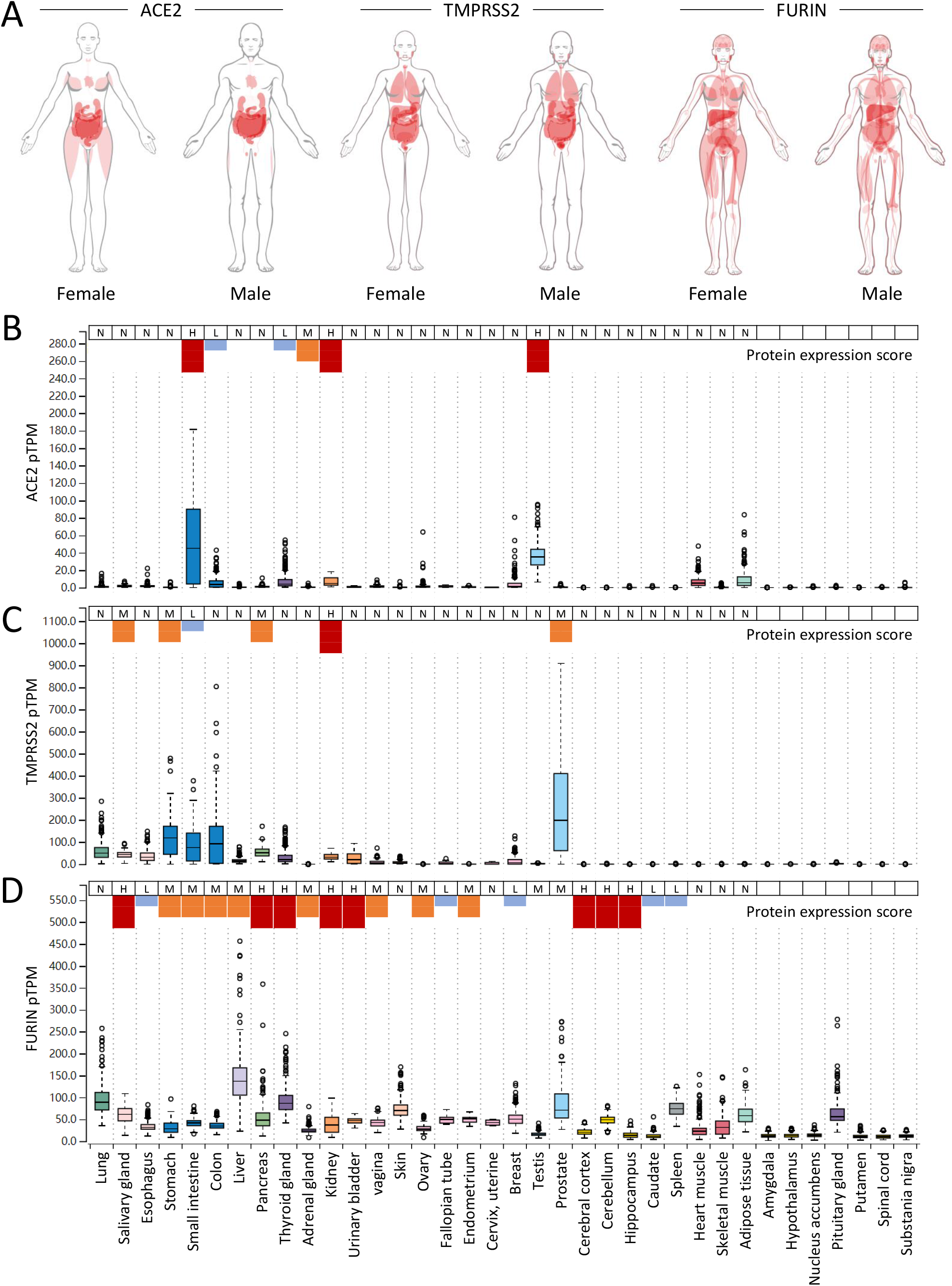
Tissue distribution of ACE2, TMPRSS2 and Furin proteases. (A) the anatomogram of the expression of *ACE2, TMPRSS2*, and *FURIN* in male and female human tissues. The colour strength corresponds to the gene expression level. (B-D) the mRNA expression level and protein expression score of ACE2 (B), TMPRSS2 (C) and Furin (D) in the manifold tissues and organs. N: negative; L: low expression shown in short blue column; M: median expression, shown in medium sized orange column; H: high expression, shown in long red column. TPM: transcripts per million; pTPM: all TPM values per sample scaled to a sum of 1 million TPM. Note: the RNA expression data were retrieved from GTEx database, the protein expression scores were retrieved from HPA database, in which the protein scores of last seven tissues are missing, therefore not indicated. Figure S1 shows RNA and protein expression on more tissues from HPA database.

The expression of TMPRSS2 also had a tissue-specific pattern, but it was different from that of ACE2 in terms of tissue distribution (Figure 1A, 1C, S1B and S1E). There were 5 tissues co-expressing relatively high mRNA levels of TMPRSS2 and ACE2, include the small intestine, colon, thyroid gland, kidney and breast. At the protein level, only the small intestine and kidney showed co-expression. Interestingly, the prostate expressed the highest level of TMPRSS2 mRNA but had an undetectable level of ACE2 protein. These results suggest that TMPRSS2 and ACE2 are not often co-expressed in the same tissues. Notably, considerable TMPRSS2 mRNA was expressed in the lung, the target tissue of SARS-CoV-2 virus, which was consistent with a promoting role of TMPRSS2 in SARS-CoV-2 infection.

As compared with ACE2 and TMPRSS2, the expression of Furin protease was much less specific at both the mRNA and protein levels, though some tissues, such as the liver and lung, did express much higher than the others. Regarding the 5 tissues co-expressing TMPRSS2 and ACE2, the thyroid gland and kidney appeared to simultaneously express high levels of Furin mRNA or protein (Figure 1A, 1D, S1C and S1F).

### Expression of ACE2, TMPRSS2 and Furin *in situ* in tissues

Tissues generally comprise multiple types of cells; therefore, the expression level of a specified gene at the tissue scale may not be representative of its level at a certain type of cell. To address this issue, we first analysed the protein expression of ACE2, TMPRSS2 and Furin *in situ* from immunohistochemistry (IHC) images. Tissues highly expressing ACE2 either at the mRNA level or protein level were included for analysis. In addition, the putative SARS-CoV-2 target tissues proposed by previous studies, irrespective of ACE2 expression level, were also included. This resulted in a list of 15 tissues in total (Figure 2, S2 and S3), which, based on the IHC intensity of ACE2, could be roughly categorised into 3 groups. The ACE2-high group contained the kidney, stomach, small intestine, colon, gallbladder and testis. In this group of tissues, ACE2 was generally expressed at the apical region, facing the luminal surface of epithelial cells (Figure 2D-E, S2D-F, and S3B-D), which means that efficient host entry of virus could only take place from the luminal side. This rule may not be applied to testis as high level of ACE2 expression was also detected in the stromal cells (Figure 2F and S2F), which are not polarized and could be potentially accessed by virus from the blood stream. The ACE2-low group contained liver, oesophagus, breast and adipose tissue, which are unlikely to be the direct targets of SARS-CoV-2 virus even though some of them showed high levels of mRNA expression, such as breast (Figure S3E) and adipose tissues (Figure S3F). There was a small amount of ACE2-positive cells in the interlobular region of liver tissue (Figure 2C and S2C), probably the cholanggiocytes as suggested below in scRNA analysis (Figure 4C). These cells might be potential SARS-CoV-2 targets under the condition that viruses come from the luminal side, which is normally rare, as the ACE2 protein was detected at the cell apical surface. The ACE2-median group contained the lung, heart, ovary, adrenal gland and thyroid gland. For lung, the major target of SARS-CoV-2 virus, there were considerable amounts of ACE2-positive cells in the alveolus lumen, which morphologically resembled macrophages and represented for the majority of ACE2-positive cells in the lung tissue (Figure 2A and S2A). All of the cardiomyocytes, and a portion of stromal cells in the ovary, adrenal gland and thyroid gland significantly expressed ACE2 (Figure 2B, 2G-H, S3G), therefore were potential virus targets in the presence of viremia.

**Figure 2.**
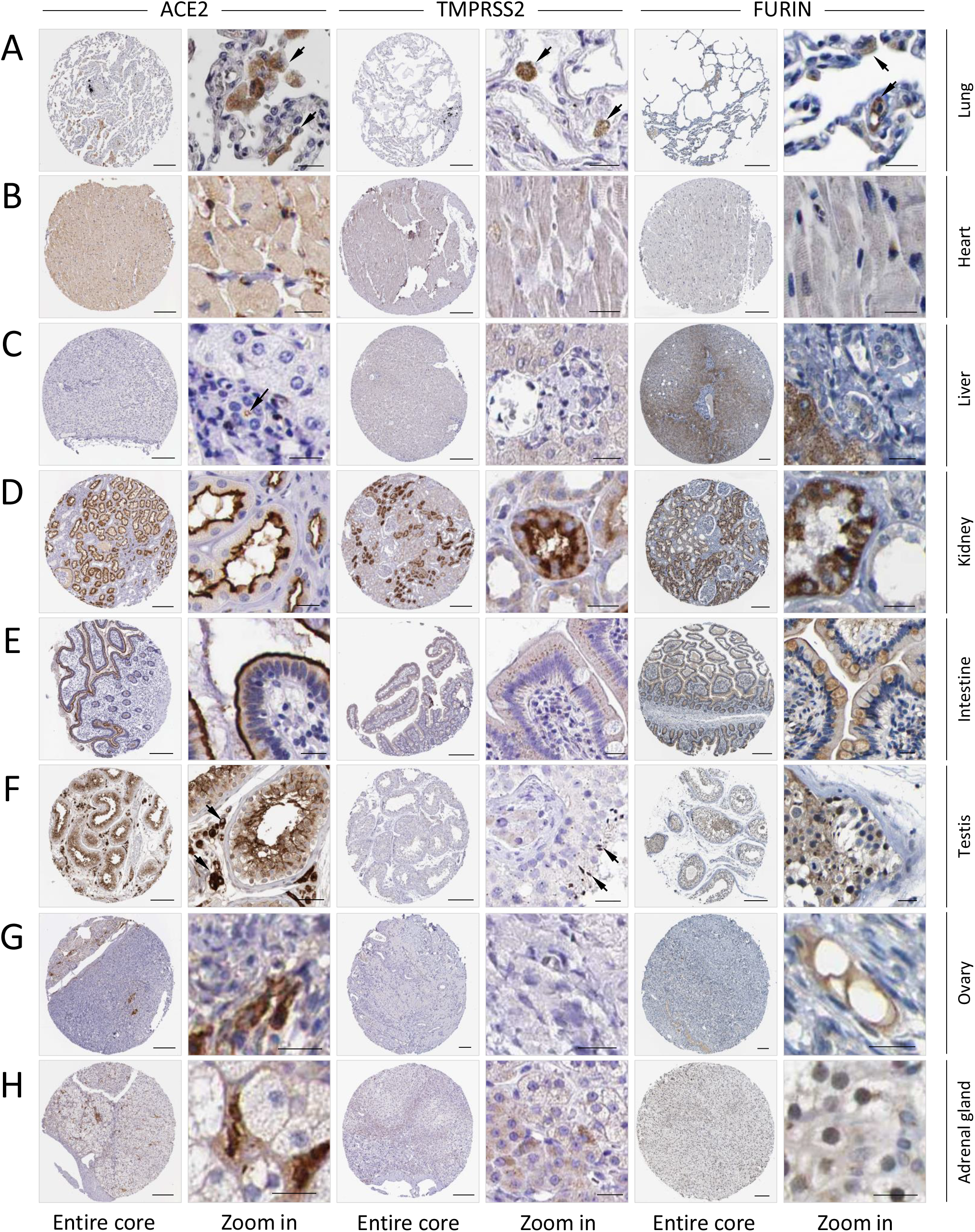
Protein expression of ACE2, TMPRSS2 and Furin *in situ* in tissues. (A-H) the IHC images for the protein expression of ACE2, TMPRSS2 and Furin in the indicated tissues/organs. F. adrenal gland: fetal adrenal gland. Scale bars: 200 μm for core images, 20 μm for zoom in images. Arrows indicate the positive IHC signals (note: not all signals were indicated). Please find in supplementary Figures for image replicates (Figure S2) and images of more tissues (Figure S3).

The expression of TMPRSS2 and Furin generally was also high in ACE2-high group tissues (Figure 2D-E, S2B-D), with the exception of the testis, where TMPRSS2 seemed to be only highly enriched in the spermatids (Figure 2F). For the ACE2-low group, TMPRSS2 and Furin were also expressed at low levels (Figure S3A, S3E-F) with the exception of the liver, where the hepatocytes expressed Furin at quite high level (Figure 2C). For the ACE2-median group, while Furin was readily detected in all the tissue cells, TMPRSS2 expression could be readily detected in lung macrophages (Figure 2A and S2A), cardiomyocytes (Figure 2B and S2B) and adrenal gland cells (Figure 2H and S2H), but not the stromal cells of ovary and thyroid gland (Figure 2G, S2G and S3G), suggesting different probabilities for them to establish efficient infection by SARS-CoV-2 virus during viremia.

Together, based on the above analysis, we propose that lung macrophages, in addition to the well-known AT2 cells, may be another direct target of SARS-CoV-2 virus. In the presence of viremia, the top vulnerable targets might be the heart and adrenal gland, and then the less likely the testis, ovary and thyroid gland. Other tissues are either unlikely direct targets or incompetent for establishing efficient infection due to lack of access to virus or low expression of helping proteases.

### Identification of cell types in tissues and organs by scRNA expression profiling

To obtain comprehensive analysis of the target cells of SARS-CoV-2, we utilized the curated public databases to perform scRNA profiling of the ACE2-high and -medium tissues, including the lung, heart, kidney, intestinal track (ileum, rectum, colon), stomach, testis, ovary, thyroid and adrenal gland. In addition, we also included the liver, the ACE2-low organ, in scRNA analysis given its important clinical implications. After quality filtering (see Material and methods), We obtained a total of 147,726 cells and annotated 79 cell types involving the respiratory, circulatory, digestive, urinary, reproductive and endocrine systems.

The lung, as the pivotal respiratory organ, is one of the target organs of SARS-CoV-2. In the lung dataset, 28819 cells from three donors passed stringent quality control and represented 11 cell types. Specifically, we identified alveolar type II (AT2) cells, alveolar type I (AT1) cells, ciliated and club cells, alveolar macrophages, dendritic cells, monocytes, fibroblasts, endothelial cells, T and NKT cells, plasma and B cells. Expression of canonical cell markers for pulmonary cell types was observed in largely non-overlapping cells, including SFTPC for AT2 cells, AGER for AT1 cells, TPPP3 for ciliated cells, SCGB3A2 for club cells, CD68 for macrophages, VWF for endothelial cells, CD3D for T and NKT cells, IGHG4 for plasma and B cells (Figure 3A). In the heart dataset, we obtained 8148 high-quality cells consisting of 5 prime cell types, including cardiomyocytes, endothelial cells, fibroblasts, smooth muscle cells (SMC) and macrophages on the basis of their respective molecular features (Figure 3B).

**Figure 3.**
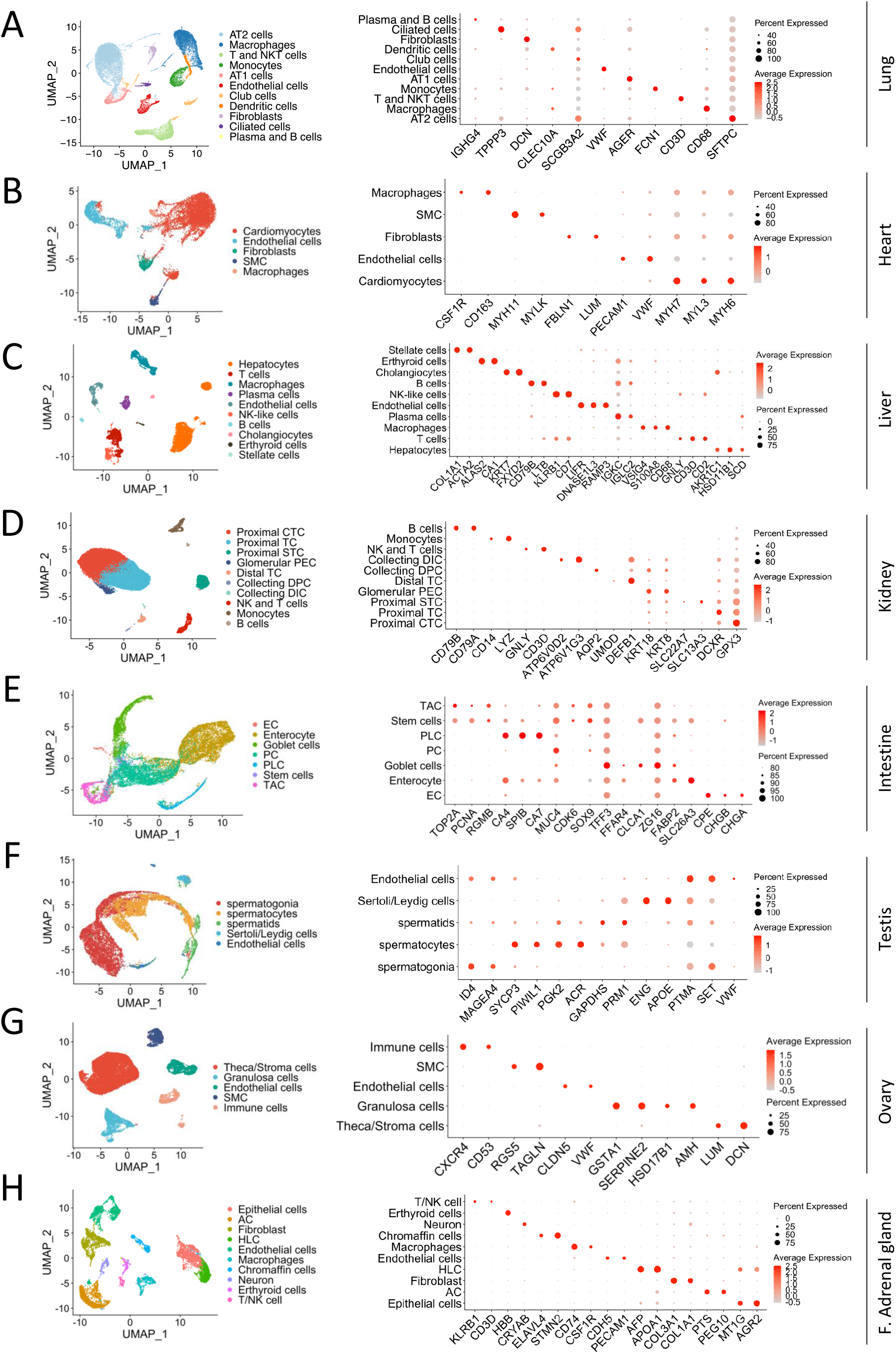
Single cell profiling of the cell types in tissues and organs. UMAP plots on the left show single cell transcriptomic profiling of the cell types from the indicated tissues/organs. Dot plots on the right represent the expression of canonical marker genes of each cell types. AT2: alveolar type II, AT1: alveolar type I, SMC: smooth muscle cells, NK: natural killer, CTC: convoluted tubule cells, TC: tubule cells, STC: straight tubule cells, PEC: parietal epithelial cells, TC: tubule cells, DPC: duct principal cells, DIC: distal tubule cells, EC: enteroendocrine cells, PC: paneth cells, PLC: paneth-like cells, TAC: transient amplifying cells, AC: adrenocortical cell, HLC: hepatocyte-like cell and F. adrenal gland: fetal adrenal gland.

As such, the cell population of hepatic tissues from 5 human livers was classified into ten cell types based on the expression of known markers, which was shown in Figure 3C. In addition, 23,366 kidney cells from 3 donors were identified and grouped into 10 cell types. The detailed cell categories of kidney and the expression of specific cell markers were shown in Figure 3D.

For integrative analysis of the human intestinal mucosa at single cell level, we pooled 14,207 intestinal cells together (6167 cells from two ileum samples, 4411 cells from two colon samples and 3629 cells from two rectum samples). In total, these cells were partitioned into 7 main cell types, containing enteroendocrine cells (EC), enterocytes, goblet cells, paneth cells (PC), paneth-like cells (PLC), stem cells and transient amplifying cells (TAC) (Figure 3E). Additionally, we also investigated the gastric mucosa dataset incorporating 5281 cells and finally annotated 9 cell types (Figure S4A-B).

To characterize the single-cell profiling of human reproductive organs, the testis and ovarian datasets were thoroughly analysed. In the testis datasets from adult human sorted spermatogonia, spermatocytes and spermatids, we identified 12,829 cells, which were classified into 5 major cell populations for downstream analysis (Figure 3F). Similarly, we exploited 27,857 cells from 5 adult women ovaries, and defined 5 major cell types, including theca cells, granulosa cells, endothelial cells, SMC and immune cells (Figure 3G).

The adrenal gland and thyroid, as the significant parts of the endocrine system, were also included into our research. We took advantage of 9809 cells from the foetal adrenal gland and 8966 cells from the thyroid to construct the single cell atlas (Figure 3H, Figure S4H). Based on the expression of specific cell markers, we found that the atlas of the fetal adrenal gland mainly comprised epithelial cells, adrenocortical cell (AC), hepatocyte-like cell (HLC), and chromaffin cells, while the atlas of the thyroid mainly consisted of thyroid follicular cells, endothelial cells and SMC (Figure 3H, Figure S4I).

### Single cell transcriptomic profiling of ACE2, TMPRSS2 and Furin proteases in distinct cell types

To determine the mRNA expression level of ACE2, TMPRSS2 and Furin genes in distinct tissue cells, we systematically explored the established cell atlases of different organs and tissues. We found the expression level of ACE2, TMPRSS2 and Furin genes varied significantly across the tissues and organs analyzed (Figure 4). Importantly, in certain tissues, these genes were only expressed in particular cell types. For example, the SARS-CoV-2 receptor ACE2 was mainly expressed in AT2 cells in the lung (Figure 4A), and cholangiocytes and hepatocytes in the human liver (Figure 4C), which was consistent with results of previous studies (Chai et al., 2020; Zhao et al., 2020b) and confirmed our above analysis on the protein expression of ACE2. We further investigated the mRNA expression of TMPRSS2 and Furin proteases and confirmed that these genes were also expressed in the liver cholangiocytes and AT2 cells at the high level (Figure 4A and C). Notably, in the adult ovary, we found that only stroma cells expressed the ACE2 and Furin genes (Figure 3G). In addition, the spermatogonia moderately expressed the ACE2 gene as well as Furin and TMPRSS2 (Figure 4F), while the Sertoli/Leydig cells expressed relatively low level of TMPRSS2, which was consistent with IHC results indicated above.

**Figure 4.**
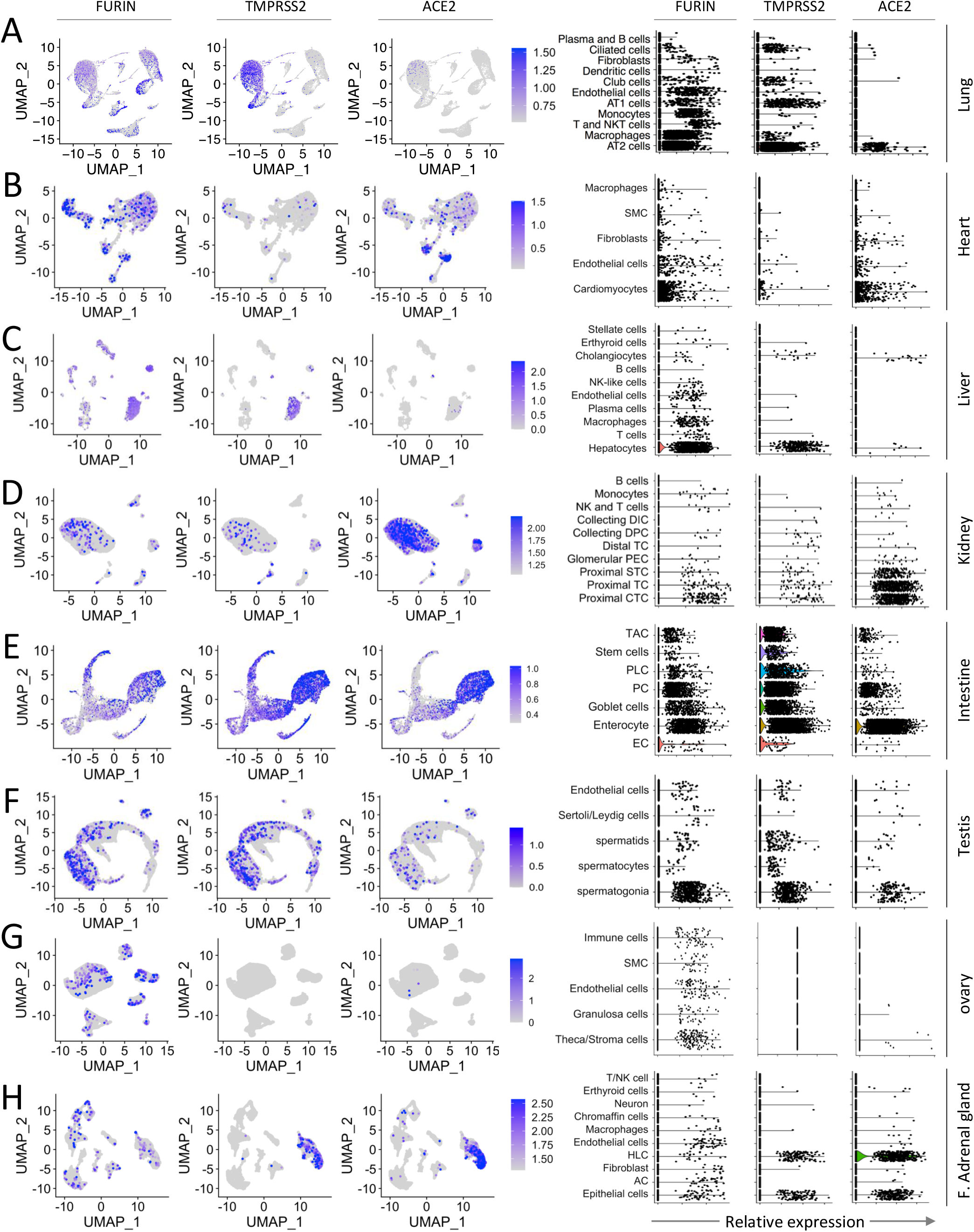
Single cell transcriptomic profiling of ACE2, TMPRSS2 and Furin proteases in distinct cell types. UMAP plots (left panel) and violin plots (right panel) show the mRNA expression of ACE2, TMPRSS2 and Furin genes in different tissue cells from the indicated tissues/organs. AT2: alveolar type II, AT1: alveolar type I, SMC: smooth muscle cells, NK: natural killer, CTC: convoluted tubule cells, TC: tubule cells, STC: straight tubule cells, PEC: parietal epithelial cells, TC: tubule cells, DPC: duct principal cells, DIC: distal tubule cells, EC: enteroendocrine cells, PC: paneth cells, PLC: paneth-like cells, TAC: transient amplifying cells, HLC: hepatocyte-like cell, AC: adrenocortical cell and F. adrenal gland: fetal adrenal gland.

Moreover, we observed that the expression of the ACE2 gene in the intestine and kidney was quite high (Figure 4D, E). Interestingly, the expression levels of the TMPRSS2 and Furin genes in the kidney were not remarkable (Figure 4D), which is inconsistent with the IHC results (Figure 2D). In addition, the high expression of the ACE2 and Furin genes were found in cardiomyocytes (Figure 4B), which fits well with the IHC results (Figure 2B). Moreover, the significant expression of the ACE2 gene was also shown in the cells of the fetal adrenal gland, specifically in HLC and epithelial cells, as well as the significant expression of the TMPRSS2 and Furin genes (Figure 4H). By contrast, in the thyroid, there was little expression of the ACE2 despite the high expression of TMPRSS2 and Furin (Figure S4J-K), which disagreed with the IHC results that ACE2 was highly expressed in stromal cells of thyroid gland (Figure S3G). These results show that the heart and fetal adrenal gland are more likely vulnerable to the SARS-CoV-2 virus via cardiomyocytes and HLC cells, respectively.

Taken together, we comprehensively analyzed the expression of ACE2, TMPRSS2 and Furin genes in different tissue cells by feat of the scRNA-seq profiling and laid the foundation for identifying the target cells of SARS-CoV-2.

### Characterization of the co-expression features of ACE2, TMPRSS2 and Furin

Considering the fact that cell receptors like ACE2, together with the proteases such as TMPRSS2 and Furin, are required for the SARS-CoV-2 virus to efficiently infect cells, it is imperative to characterize the co-expression patterns between ACE2, TMPRSS2 and Furin. Accordingly, we resorted to single-cell transcriptomes to profile their co-expression patterns in the cell types indicated.

The results showed that the co-expression feature of these three genes were quite dominant in the intestine, specifically in enterocytes (Figure 5E), and the proportion of ACE2 positive cells expressing either or both TMPRSS2 and Furin in the intestine is higher than the other ACE2-high tissues such as the kidney (Figure 5D), testis (Figure 5F) and stomach (Figure S4E-G). In detail, as shown in Figure 5D, though the cells in the kidney expressed ACE2, TMPRSS2 and Furin at moderate to high levels, there was only a small proportion of ACE2 positive cells expressing either or both TMPRSS2 and Furin. The ACE2-low liver tissue displayed an interesting expression pattern, showing that the co-expression feature of ACE2 with Furin and TMPRSS2 in the hepatocytes and cholangiocytes was remarkable (Figure 5C), but the basal expression level with ACE2 in both RNA and protein level was much lower (Figure 1B and 4C), arguing against the liver as a primary SARS-CoV-2 target.

**Figure 5.**
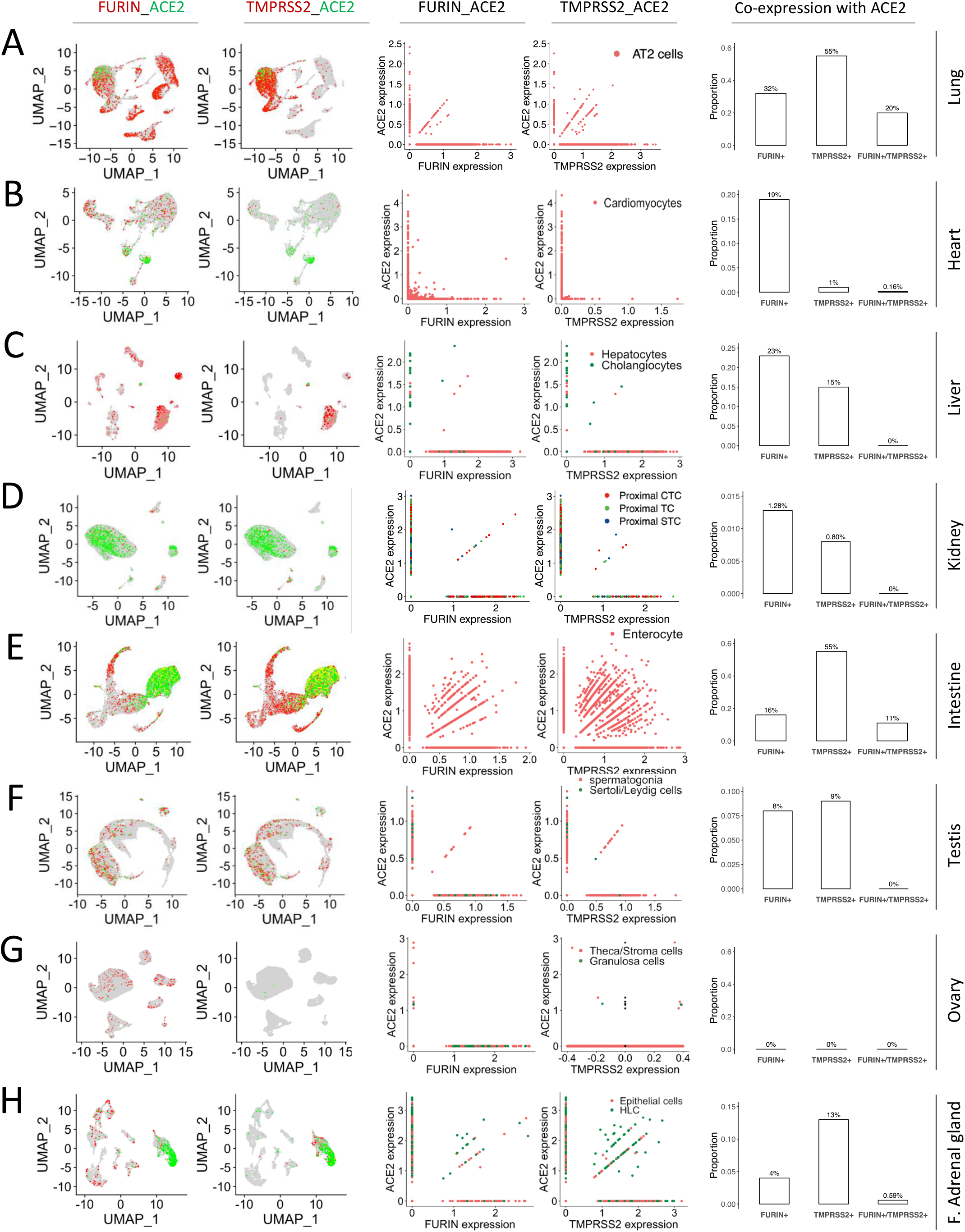
Characterization of the co-expression patterns of ACE2, TMPRSS2 and Furin at single cell level. UMAP plots (left panels) show the co-expression of ACE2 with Furin or TMPRSS2 in the indicated tissues/organs. The different colors in UMAP plots are corresponding to different genes as indicated on the top of each graph column. Scatter plots (two middle panels) illustrate the expression correlation of ACE2 with Furin or TMPRSS2 in the AT2 cells (A), cardiac cardiomyocytes (B), liver hepatocytes and cholangiocytes (C), the proximal tubule cells of kidney (D), the enterocyte of intestinal tract (E), the spermatogonia and Sertoli/Leydig cells in the testis (F), the stroma cells and granulosa cells in the ovary (G), the epithelial cell and HLC in the fetal adrenal gland (H). The colors in scatter plots indicate the corresponding cell types depicted on the up-right corner. Barplots (right panels) show the proportion of ACE2 positive cells expressing either or both FURIN and TMPRSS2. The number above the bar in the barplots indicating the corresponding percentage. CTC: convoluted tubule cells, TC: tubule cells, STC: straight tubule cells, HLC: hepatocyte-like cell, and F. adrenal gland: fetal adrenal gland.

In the ACE2-medium tissues, we found that the lung displayed the co-expression features, especially in AT2 cells, and a highest percentage of ACE2 positive cells expressing either or both TMPRSS2 and Furin in the investigated organs and tissues (Figure 5A), which provided the evidence that SARS-CoV-2 posed the serious threat to the lung. Moreover, we observed a significant co-expression of the ACE2 gene with TMPRSS2 and Furin in the epithelial cells and HLC of the fetal adrenal gland (Figure 5H), manifesting that the fetal adrenal gland may be a target of SARS-CoV-2. Furthermoreconsistent with the results above, a portion of cardiomyocytes simultaneously expressed ACE2 and Furin (Figure 5B). However, few cells co-expressed ACE2, TMPRSS2 and Furin simultaneously in the ovary (Figure 5G) and thyroid gland (Figure S4M-N).

### The stomach is a barrier against SARS-CoV-2 virus

Since the normal stomach is highly acidic, it may conceivably serve as a physical barrier against SARS-Co-V-2 virus. In order to test this idea, we made viruses pseudotyped with SARS-CoV-2 spike protein. The same amounts of pseudo-viruses were pretreated under acidic conditions with pH of 1.0, 2.0, 4.0, or 7.0, respectively, and then used to infect 293T-ACE2 cells and Hela-ACE2 cells. As shown in Figure 6, the viruses were completely inactivated and incapable of infecting both types of cells anymore under the pH of 1.0 and 2.0, a condition resembling the normal acidity of fasting stomach (Weiss and Clark, 1985). Even under the pH of 4.0, the viral infectivity was significantly compromised to 23% – 52% of normal level. Thus, the SARS-CoV-2 virus was likely acid-instable, which is consistent with clinically uncommon symptom in gastrointestinal tract.

**Figure 6.**
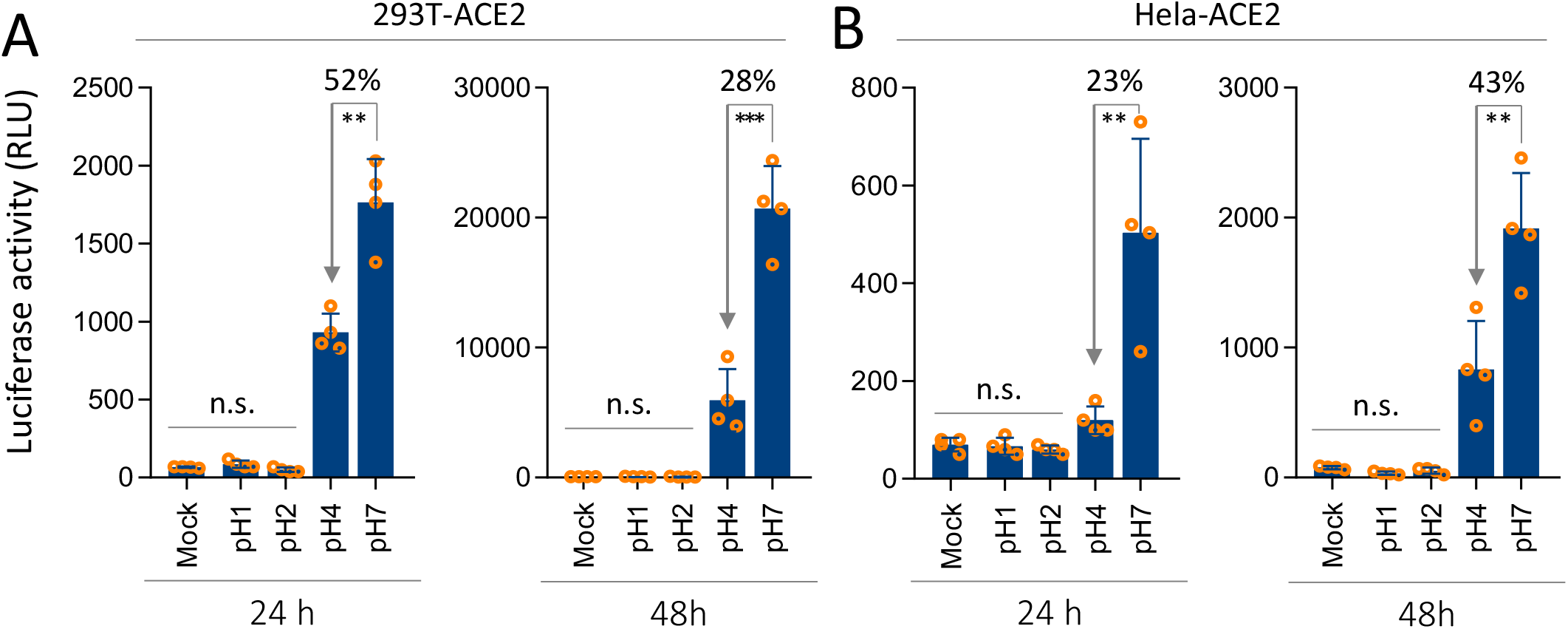
Compromised infectivity of SARS-CoV-2 pseudo-virus in acidic environment. (A, B) the effects of low pH on the activities of SARS-CoV-2 pseudo-virus to infect 293T-ACE2 cells (A), or Hela-ACE2 cells (B), as determined by luciferase assay 24 hours (left) and 48 hours (right) post infection. n.s.: not significant; **: *p* < 0.01; ***: *p* < 0.001

## DISCUSSION

Previous analysis based solely on the mRNA expression of ACE2 suggested that multiple tissue cells may be potential targets of the novel coronavirus SARS-CoV-2, including cells from almost every important human system. However, the COVID-19 patients primarily display symptoms in the respiratory system, where ACE2 is expressed in only a small portion of cells. Even for severely ill patients, the injuries or symptoms in the ACE2-high organs such as the kidney and intestinal tract are relatively uncommon, with a rate of 1%-4.3% for acute kidney injury (Guan et al., 2020; Wang et al., 2020a), and a rate of 3.5%-5.8% for intestinal symptoms (Guan et al., 2020; Special Expert Group for Control of the Epidemic of Novel Coronavirus Pneumonia of the Chinese Preventive Medicine, 2020). Instead, the injuries in ACE2-median organs such as the heart are more common(12%-19%) (Guan et al., 2020; Special Expert Group for Control of the Epidemic of Novel Coronavirus Pneumonia of the Chinese Preventive Medicine, 2020). This obvious discrepancy suggests that mechanisms other than ACE2 expression level also play important roles in establishing efficient and successful infection. Based on our analysis by pscRNA expression profiling, we propose that three more factors, in addition to the mRNA level, should be taken into account for predicting the vulnerability of a specified target tissue/organ to SARS-CoV2 infection.

First, the information on the protein expression level and protein subcellular localization may influence virus infection. As shown above, the mRNA level did not often correspond to protein level, for example, the ACE2 expression in lung macrophages (Figure 2A and 4A). While ACE2 is readily detected in macrophages in IHC staining (Figure 2A), there are only several macrophages positive in ACE2 (Figure 4A). This obvious discrepancy may be resulted from limited scRNA samples analysed, which may not be the case as additional lung scRNA analysis gave results similar to that in Figure 4A (data not shown); or protein-mRNA expression inconsistency might be accountable, which actually necessitates the protein-proofed scRNA profiling analysis that incorporates information from protein expression, RNA expression and biological experimentations as well. In light of this point, the final rank list of SARS-CoV-2 vulnerable cells would be a result of combined consideration. Actually, several recent works (Bost et al., 2020; Zhou et al., 2020c) indicated that macrophages are indeed one major target cell of SARS-CoV-2, which is consistent with our analysis. Therefore, it is necessary to not only analyse mRNA expression, but also detect protein expression *in situ*. Moreover, membrane proteins such as ACE2 could be expressed either all around the surface of non-polarized stromal cells such as Leydig cells in the testis (Figure 2F and 3F), or specifically on the apical region of polarized epithelial cells such as the enterocytes in the intestinal tract (Figure 2E). This means much for virus infection as it determines how the viruses may get access to their target cells and enter. For the polarized cells like enterocytes, virus could only successfully make infection from the luminal side, where ACE2 is expressed on the apical surface but not the basal-lateral surfaces. Whereas, it was reported that coronaviruses, such as MERS-CoV, lost infectivity in highly acidic gastric fluid (Zhou et al., 2017), thus, the likelihood that SARS-CoV-2 get access to enterocytes via stomach would be low. While for the non-polarized stromal cells, virus may readily come to see them from the blood stream.

Second, the co-expression of infection co-factors determines the efficiency of successful infection. As for SARS-CoV-2, TMPRSS2 and Furin were demonstrated to be important proteases that cleave the S protein to promote host entry (Hoffmann et al., 2020; Meng et al., 2020; Walls et al., 2020). Therefore, their co-expression with ACE2, the cellular receptor for SARS-CoV-2, may dictate the vulnerability of the target tissues. Actually, we found that some ACE2-high cells, such as stromal cells in the testis and ovary, expressed TMPRSS2 at quite low levels (Figure 2F-G), suggesting that they may not be SARS-CoV-2 targets as susceptible as those co-expressing both ACE2 and TMPRSS2/Furin proteases, such as cardiomyocytes in the heart, though cardiomyocytes expressed relatively lower level of ACE2. It is conceivable that cells highly co-expressing ACE2, TMPRSS2 and Furin proteases, such as lung macrophages and stromal cells in adrenal gland, would be readily vulnerable to SARS-CoV-2 attack in the presence of viruses.

Third, the feasible routes whereby viruses gain access to their target cells are also accountable for clinical manifestations. In theory, there are two major routes for virus transmission: 1) direct entry to the luminal tracks via open entries of the body. The potentially affected tissues/organs include those from the respiratory system, digestive system and urinary tract, among which the respiratory system is much easier than the rest two to be infected, as it is almost a completely open system, while the rest two are gated by multiple means. For example, the stomach, which is a highly acidic, may serve as an effective barrier for SARS-CoV-2 to enter the intestinal track by inactivating them (Figure 6). This may explain the prominent symptoms in the respiratory system but not in the digestive and urinary systems despite high levels of co-expression of ACE2, TMPRSS2 and Furin in epithelial cells lining along the tracts of the later two systems; 2) transmission via the blood stream to the entire body. This may theoretically affect all the internal organs with stromal cells expressing ACE2, with exception for those polarized cells in which the ACE2 protein is only expressed on the apical surface unreachable by the viruses from viremia. The target tissue cells affected by this route may include cardiomyocytes, stromal cells in adrenal gland, Leydig cells in the testis, and stromal cells in the ovary and thyroid gland. Among them, only the former two, but not the later three, are more likely the true or susceptible targets of SARS-CoV-2 when considering the co-expression of helping proteases with ACE2. It should be noted that the precondition for this route is viremia, which, however to our best knowledge, was not clearly documented in COVID-19 patients up to date. Nevertheless, frequent heart injuries in COVID-19 patients (Guan et al., 2020; Zhou et al., 2020a) are consistent with the presence of occasional viremia. Under such circumstances, the adrenal gland may be another vulnerable target of SARS-CoV-2, which, to our best knowledge, is identified for the first time in this study.

Intriguingly, a recent work by Ziegler *et al* (Ziegler et al., 2020) demonstrated that ACE2 is an interferon-stimulated gene, and the immune response stimulated by SARS-CoV-2 infection resulted in upregulated expression of cytokines such as interferon, which subsequently upregulates ACE2 expression in the neighbouring cells promoting virus dissemination. Thus, in addition to the basal ACE2-TMPRSS2 co-expression as analysed in this study, which mediates the initial viral infection, a positive-feedback loop between virus infection and interferon signalling would facilitate viral dissemination by increasing ACE2 expression. This finding was further confirmed by two independent studies by Smith J *et al* (Smith et al., 2020) and Wang et al (Wang and Cheng, 2020). This factor should also be taken into account when analysing tissue injuries and clinical symptoms of COVID-19 patients.

In summary, we propose that the pscRNA profiling is a feasible way for gene expression analysis at both protein and mRNA levels. Through a systemic analysis of 36 human tissues/organs by pscRNA profiling of ACE2, TMPRSS2 and Furin proteases, we propose a rank list of tissue cells potentially vulnerable to SARS-CoV-2 attack. For initial infection, ACE2-expressing cells, with the co-expression of TMPRSS2 and Furin as a plus, in the respiratory track are the primary targets. These cells include the known lung AT2 cells, and macrophages in this study, and also the nasal epithelial cells as reported recently (WU et al., 2020a). And the likelihood of epithelial cells in digestive and urinary systems as the primary targets is low. During a period of viremia, the top internal organ targets would be cardiomyocytes, and stromal cells in the adrenal gland, as both of these express ACE2, TMPRSS2 and Furin proteases; the descent targets may be Leydig cells in the testis, and stromal cells in the ovary and thyroid gland, as these cells are un-polarized and ACE2-positve, but do not show ideal co-expression of TMPRSS2 or Furin proteases. Notably, the identification of the adrenal gland as a SARS-CoV-2 target may be quite informative for clinical practise if experimentally confirmed, because the COVID-19 disease frequently proceeds to a severe or very severe stage in a short period, which suggests systemic conditions occur probably due to deregulated endocrine systems involving the adrenal gland. This issue remains to be validated experimentally and clinically.

## DATA AVAILABILITY

Availability of supporting data: GSE122960 (http://www.ncbi.nlm.nih.gov/geo/query/acc.cgi?acc=GSE122960), GSE109816 (http://www.ncbi.nlm.nih.gov/geo/query/acc.cgi?acc=GSE109816), GSE131685

(http://www.ncbi.nlm.nih.gov/geo/query/acc.cgi?acc=GSE131685), GSE125970

(http://www.ncbi.nlm.nih.gov/geo/query/acc.cgi?acc=GSE125970), GSE134520

(http://www.ncbi.nlm.nih.gov/geo/query/acc.cgi?acc=GSE134520), GSE109037

(http://www.ncbi.nlm.nih.gov/geo/query/acc.cgi?acc=GSE109037), GSE134355

(http://www.ncbi.nlm.nih.gov/geo/query/acc.cgi?acc=GSE134355), GSE118127

(http://www.ncbi.nlm.nih.gov/geo/query/acc.cgi?acc=GSE118127).

## ACKNOWLEDGEMENT

We thank Dr. Yan Zhang from the institute of biotechnology, Beijing, for assisting database analysis.

## SUPPLEMENTAL FIGURES LEGENDS

**Figure S1.**
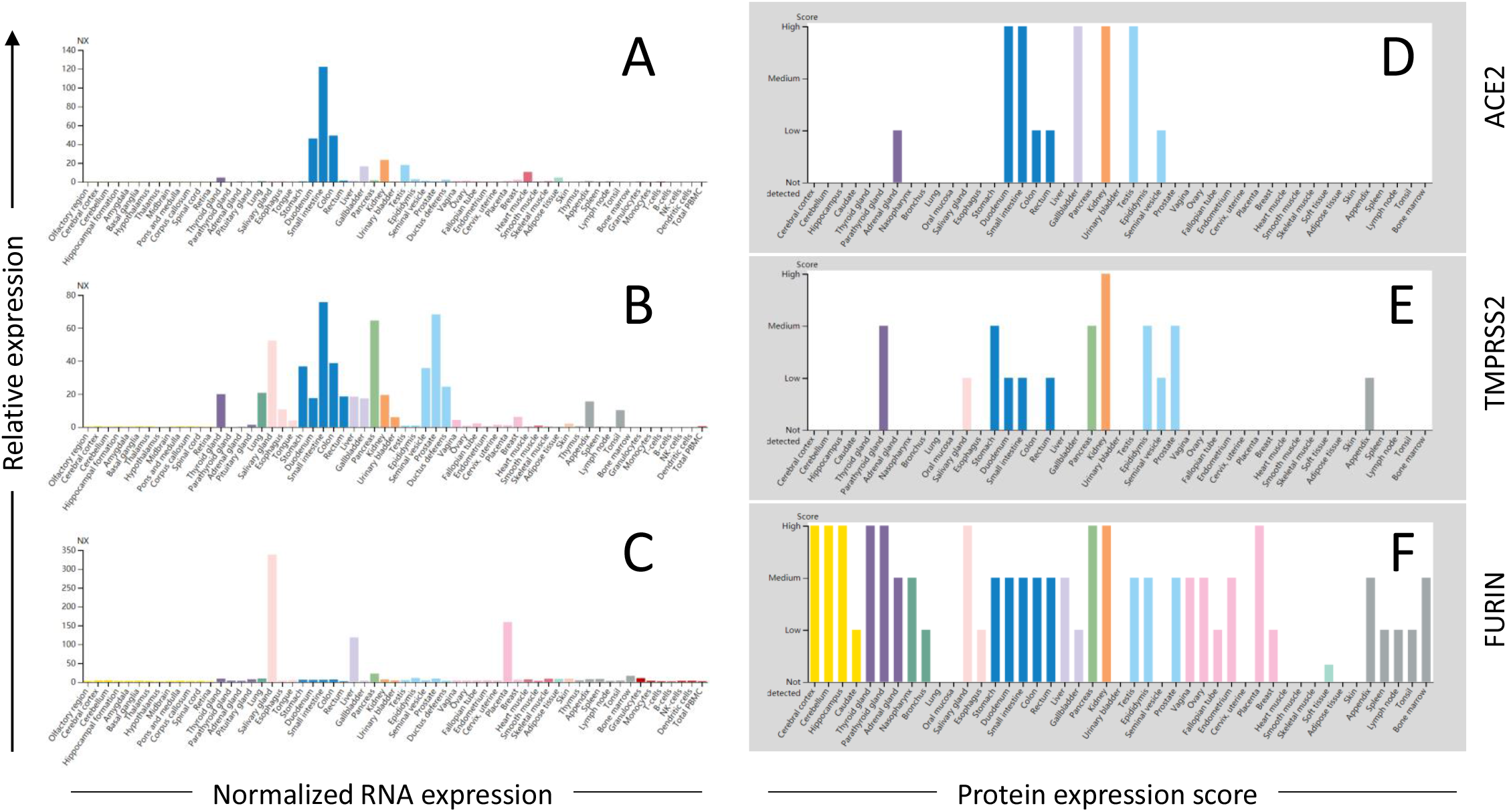
Tissue distribution of ACE2, TMPRSS2 and Furin proteases. (A-C) The normalized mRNA expression of ACE2, TMPRSS2 and Furin in 61 human tissues and organs from HPA database. (D-F) The protein expression scores for ACE2, TMPRSS2 and Furin in 44 human tissues and organs from HPA database. Note: Figure S1, derived directly from HPA database, is an extension of the data shown in Figure 1, generally with expression information from more tissues.

**Figure S2.**
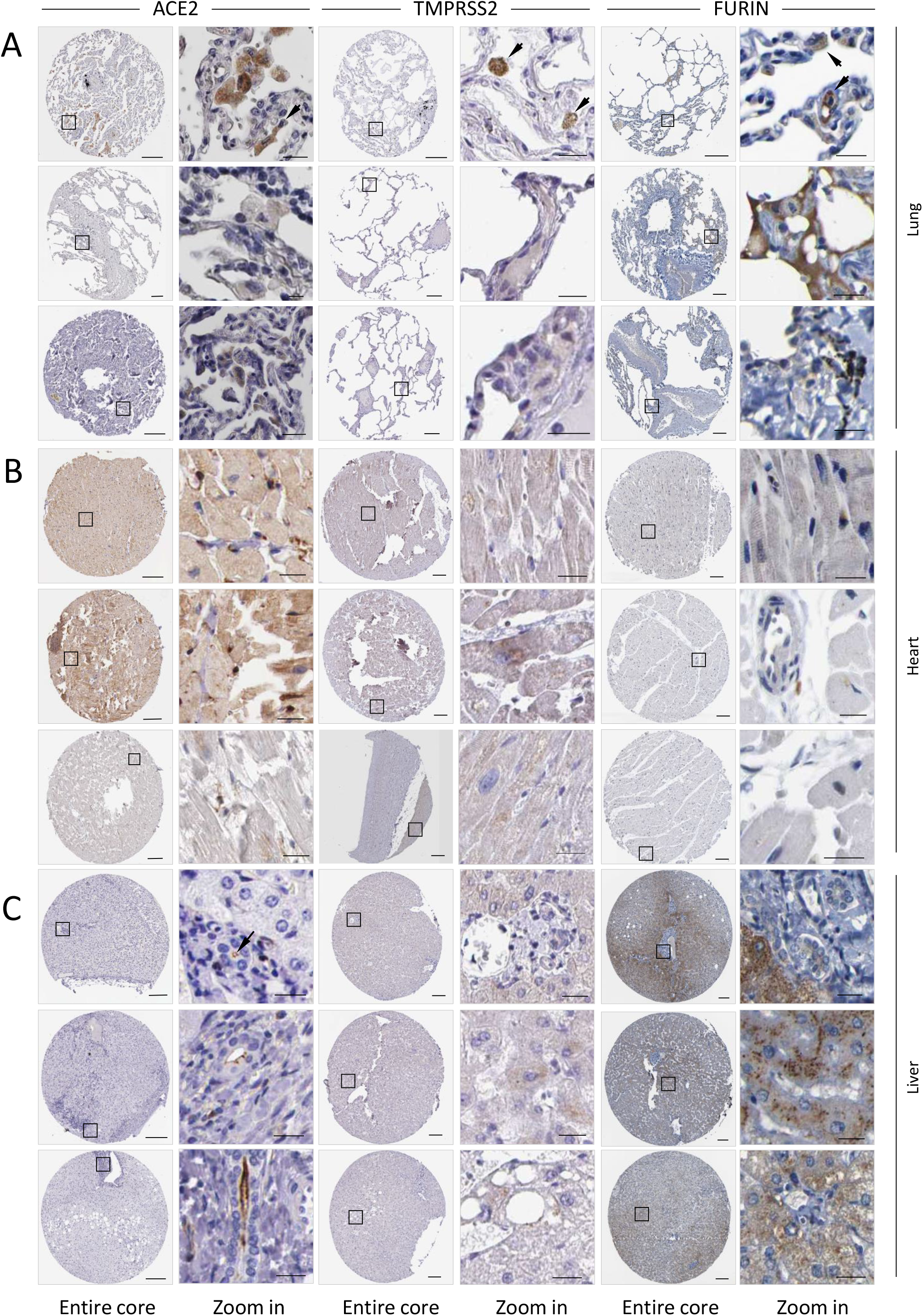
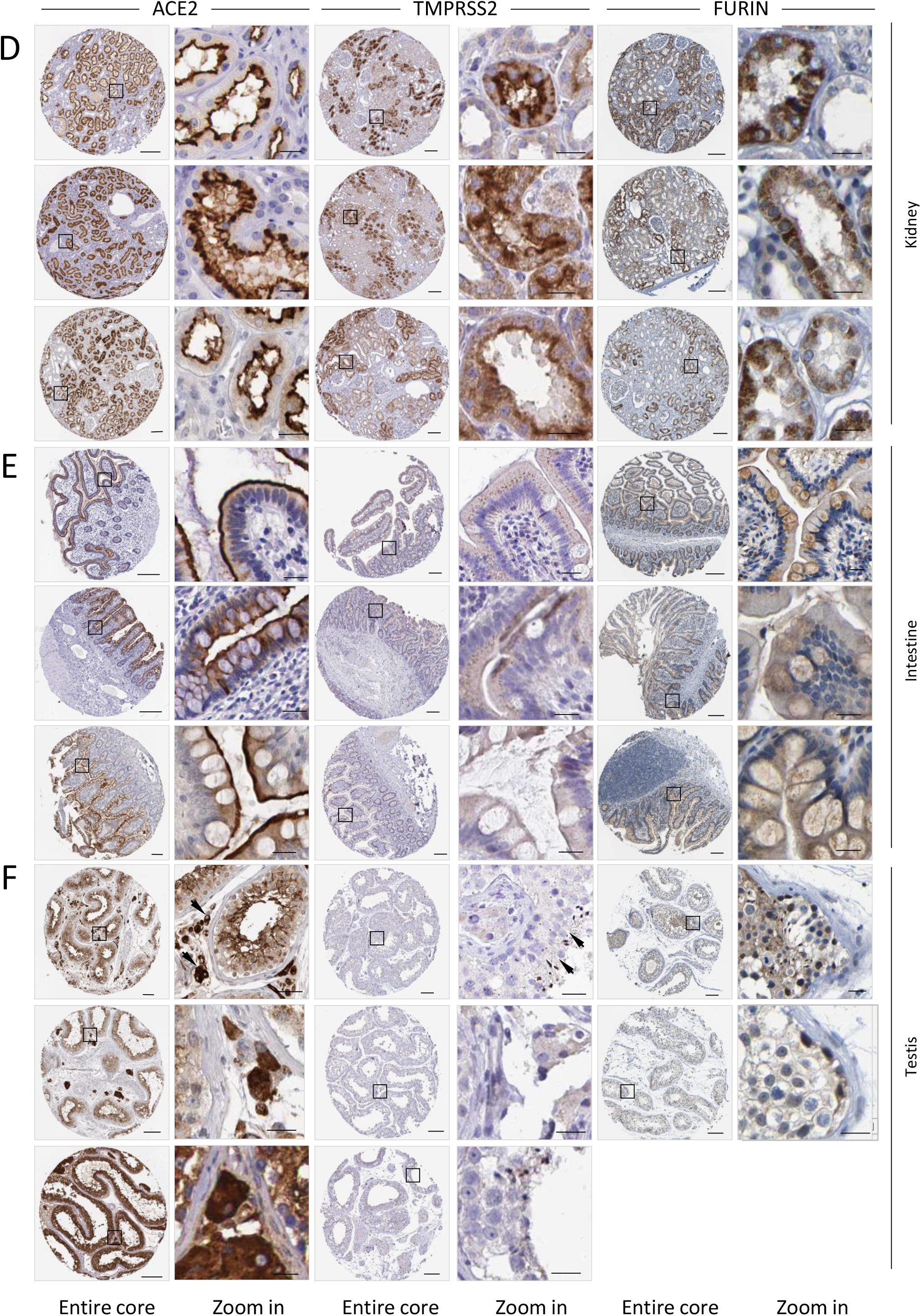
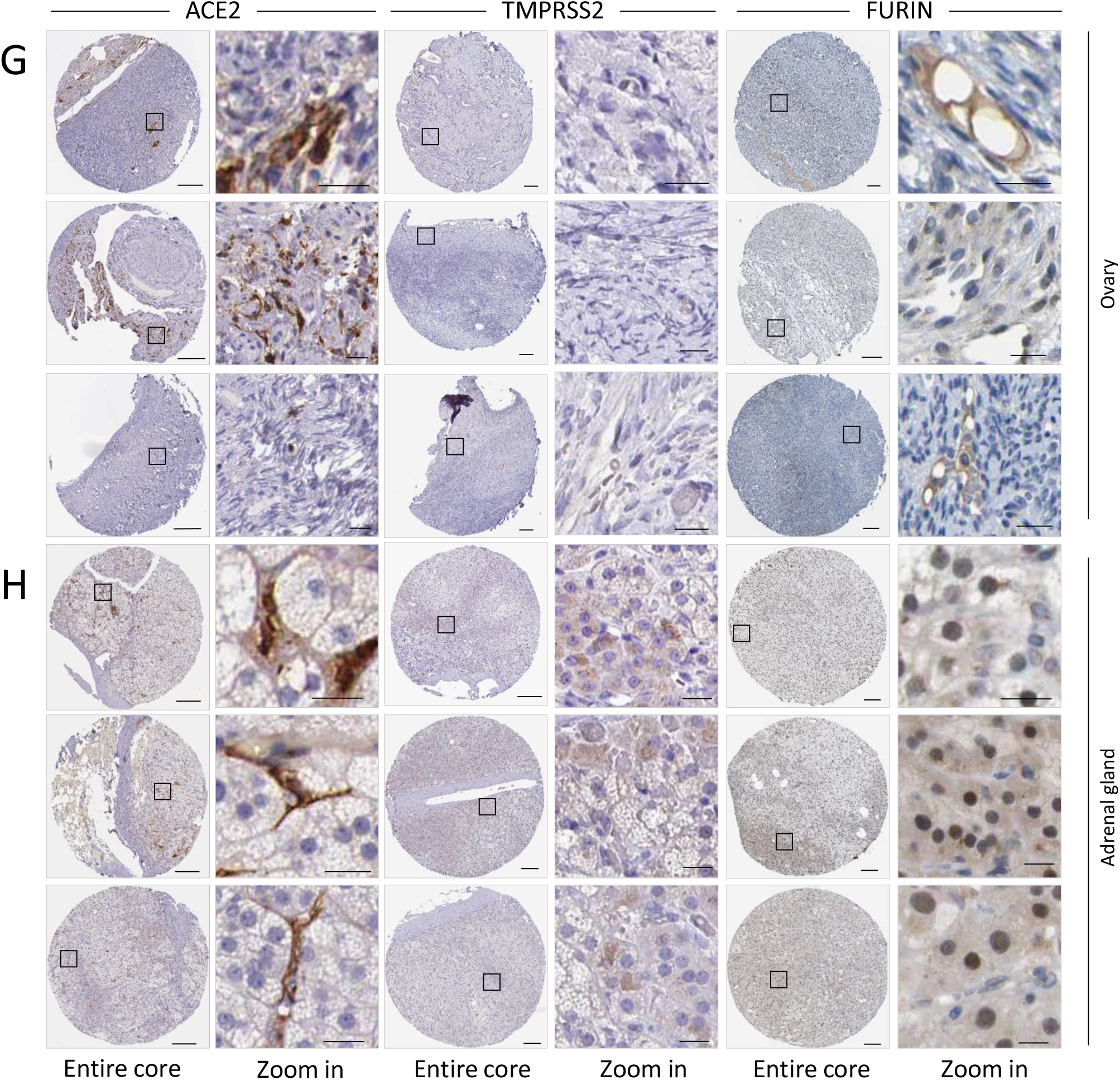
Replicate images for protein expression of ACE2, TMPRSS2 and Furin *in situ* in tissues shown in Figure 2. Generally, 3 IHC images were shown for each tissue, except for Furin expression in testis, where only 2 images are available in the HPA database. Scale bars: 200 μm for core images and 20 μm for zoom in images.

**Figure S3.**
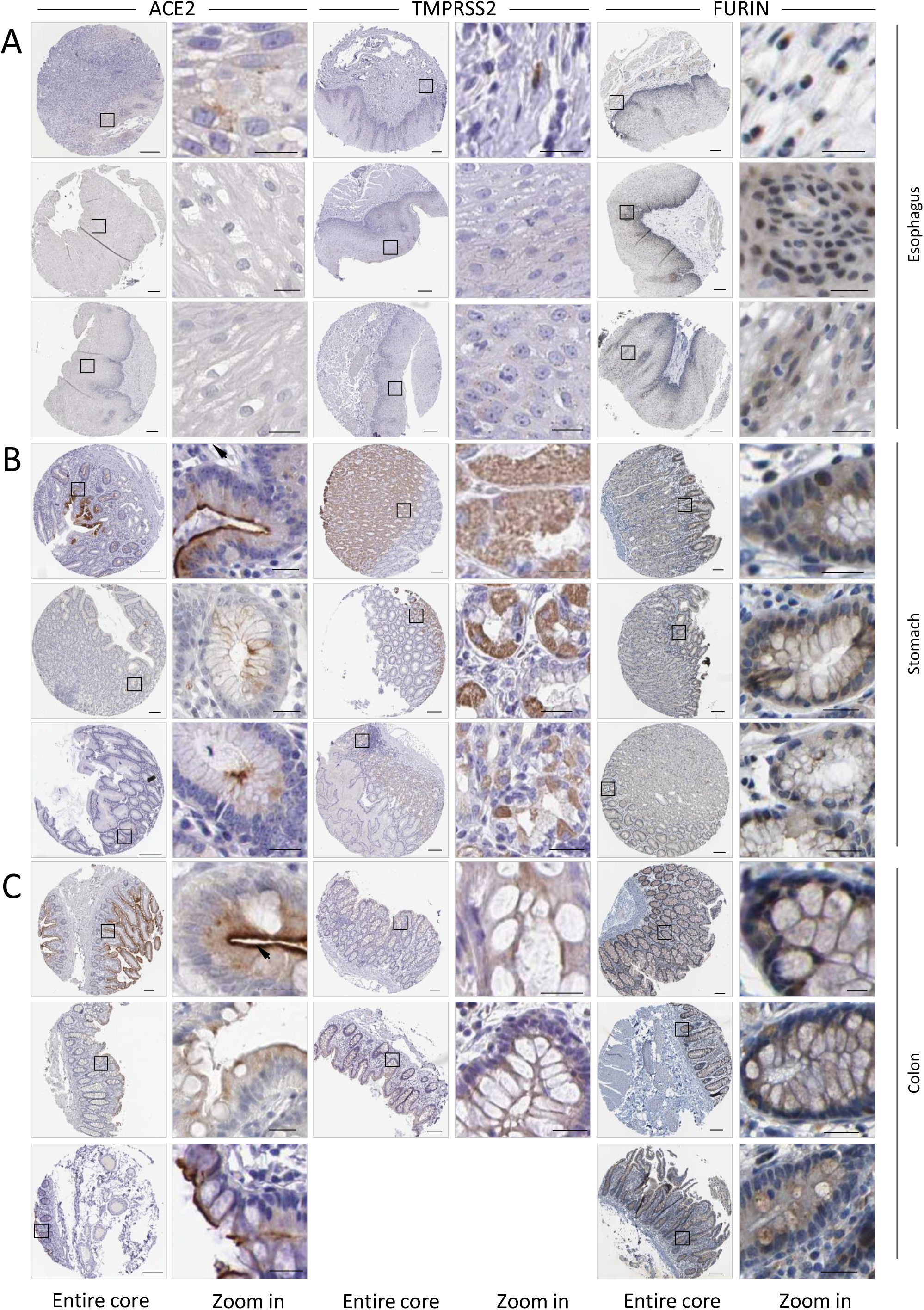
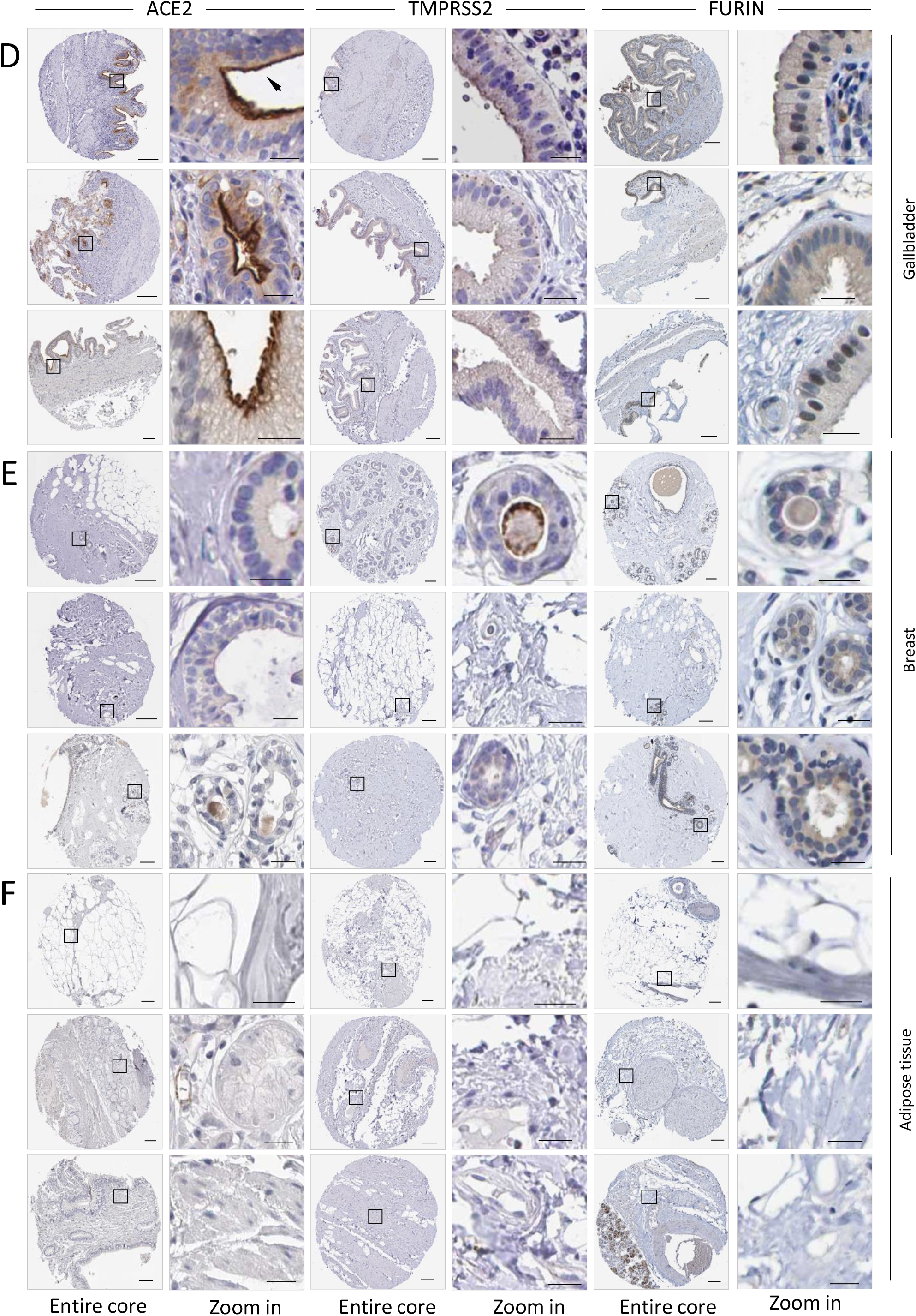
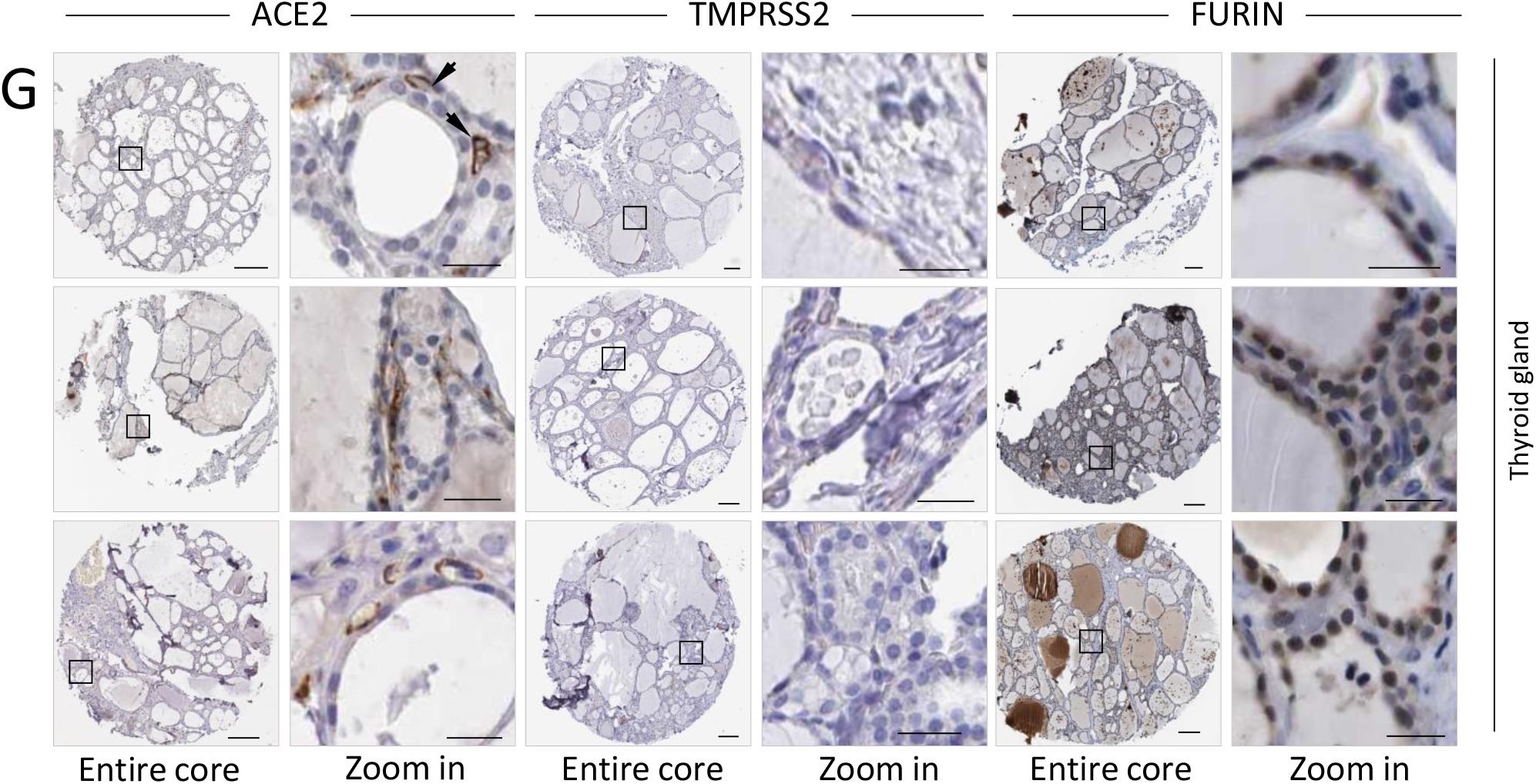
Protein expression of ACE2, TMPRSS2 and Furin *in situ* in tissues other than those indicated in Figure 2. The IHC images for ACE2, TMPRSS2 and Furin in tissues of the oesophagus (A), stomach (B), colon (C), gallbladder (D), breast (E), adipose tissue (F), and thyroid gland (G). Scale bars: 200 μm for core images and 20 μm for zoom in images. Arrows indicate the positive IHC signals.

**Figure S4.**
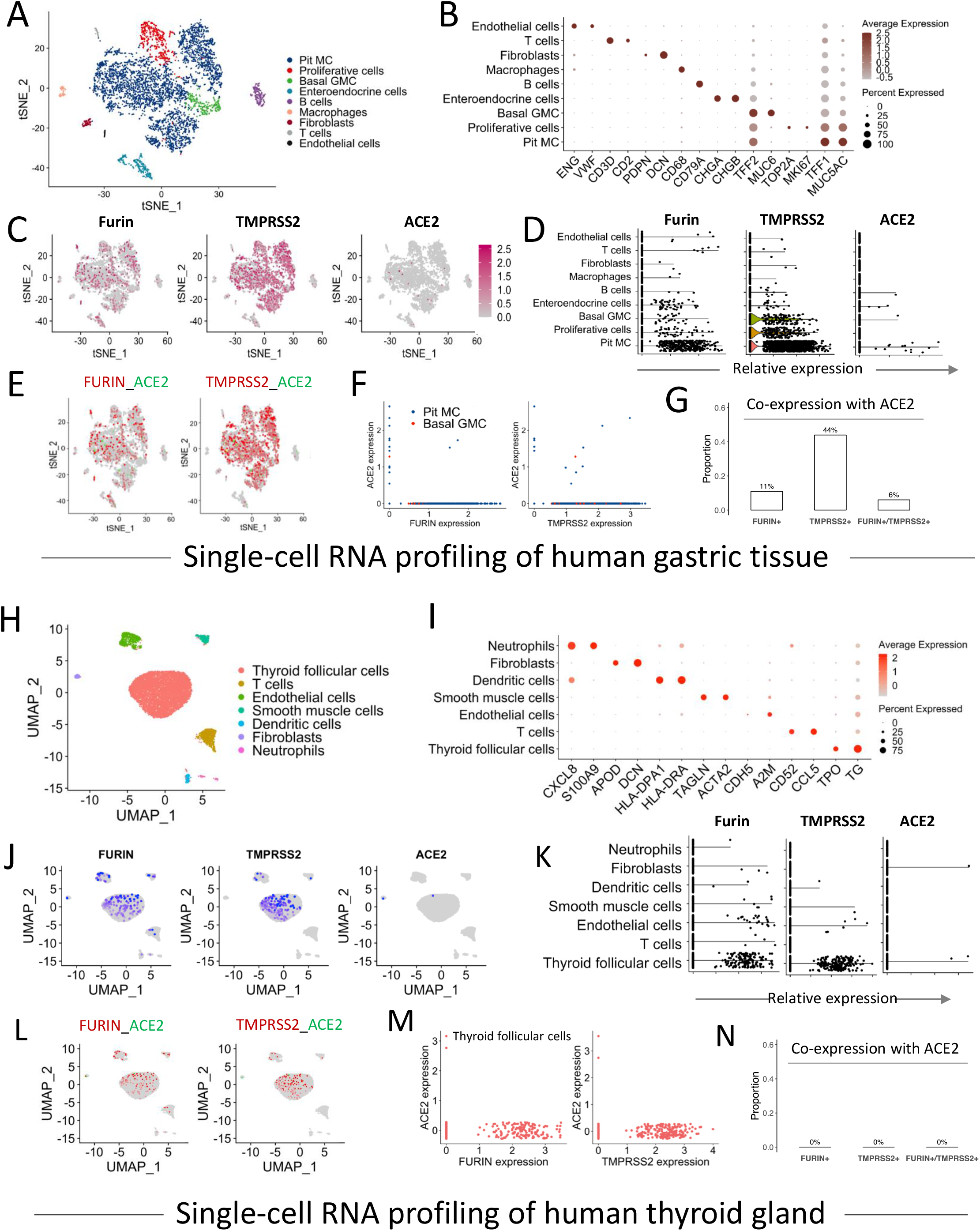
Single cell transcriptomic profiling of ACE2, TMPRSS2 and Furin expression in the gastric tissue and the thyroid gland. (A) tSNE plot showing different cell types in the stomach. (B) Dot plot showing the expression level of specific cell markers in each cell types of the gastric tissue. (C) tSNE plot revealing the expression distribution of Furin, TMPRSS2 and ACE2 in the stomach. (D) Violin plot showing the expression level of Furin, TMPRSS2 and ACE2 in different cell types of the stomach. (E-F) Characterization of the co-expression feature of ACE2, Furin and TMPRSS2 in the stomach. (G) Barplot shows the proportion of ACE2 positive cells expressing either or both FURIN and TMPRSS2 in the stomach. (H) UMAP plot representing different cell types in the thyroid gland. (I) Dot plot showing the expression level of known cell markers in each cell type of the thyroid gland. (J) UMAP plot illustrating the expression distribution of Furin, TMPRSS2 and ACE2 in the thyroid gland. (K) Violin plot showing the expression level of Furin, TMPRSS2 and ACE2 in different cell types of the thyroid gland. (L-M) Characterization of the co-expression feature of ACE2, Furin and TMPRSS2 in the thyroid gland. (N) Barplot shows the proportion of ACE2 positive cells expressing either or both FURIN and TMPRSS2 in the thyroid gland. MC: mucous cells and GMC: gland mucous cells.

**Supplementary Table 1.** Summary of sample information for single-cell analysis

| Organs/tissues | GEO accession NO. | Number of samples | Single cell sequencing technology | Number of cells | Mean counts per cell |
| --- | --- | --- | --- | --- | --- |
| lung | GSE122960 | 3 | 10x Genomics | 28819 | 5421 |
| heart | GSE109816 | 12 | SMART-seq2 | 8148 | 69059 |
| liver | GSE115469/<br>Human Cell Atlas<br>Data Portal | 5 | 10x Genomics | 8444 | 2190 |
| kidney | GSE131685 | 3 | 10x Genomics | 23366 | 2270 |
| intestine | GSE125970 | 6 (ileum:2;colon:2;rectum:2) | 10x Genomics | 14207 | 13761 |
| stomach | GSE134520 | 3 | 10x Genomics | 5281 | 3249 |
| testis | GSE109037 | 4 | 10x Genomics | 12829 | 15174 |
| ovary | GSE118127 | 10 | 10x Genomics | 27857 | 1990 |
| fetal adrenal gland | GSE134355 | 1 | Mirowell-seq | 9809 | 1495 |
| thyroid | GSE134355 | 1 | Mirowell-seq | 8966 | 880 |

**Supplementary Table 2.** Summary of the databases, software and algorithms

| RESOURCE | SOURCE |
| --- | --- |
| **Databases** |  |
| Human Protein Atlas | http://www.proteinatlas.org |
| The Genotype-Tissue Expression(GTEx) database | https://m.gtexportal.org |
| Gene Expression Omnibus (GEO) database | https://www.ncbi.nlm.nih.gov/geo |
| Human Cell Atlas Data Portal | https://data.humancellatlas.org/ |
| **Software and Algorithms** |  |
| Seurat v3.1.4 | https://github.com/satijalab/seurat |
| Harmony v1.0 | https://github.com/immunogenomics/harmony |
| clusterProfiler v3.14.3 | https://bioconductor.org/packages/release/bioc/html/clusterProfiler |
| R v3.6.1 | https://www.r-project.org/ |

**Supplementary Table 3.** recipe for pH calibration

| pH | Pseudo-virus | HCL (0.5 mol/L) | NaOH (2 mol/ L) | DMEM |
| --- | --- | --- | --- | --- |
| 1.0 | 200 µl | 60 µl | 42 µl | 98 µl |
| 2.0 | 200 µl | 40 µl | 28 µl | 132 µl |
| 4.0 | 200 µl | 10 µl | 7 µl | 183 µl |
| 7.0 | 200 µl | 0 µl | 0 µl | 200 µl |

